# Molecular dynamics simulations reveal molecular mechanisms for the gain and loss of function effects of four *SCN2A* variants

**DOI:** 10.1101/2024.02.19.580930

**Authors:** Nisha Bhattarai, Ludovica Montanucci, Tobias Brünger, Eduardo Pérez-Palma, William Martin, Iris Nira Smith, Charis Eng, Feixiong Cheng, Ingo Helbig, Rikke S Møller, Andreas Brunklaus, Stephanie Schorge, Dennis Lal

**Author notes:** Department of Physiology and Biophysics, Case Western Reserve University, School of Medicine, Cleveland, OH, 44106, USA. Center for Neurogenetics, Department of Neurosciences, University of Texas Houston, TX.

## Abstract

*SCN2A* gene disorders cover a wide range of medical conditions, from epileptic encephalopathies to neurodevelopmental disorders. The variants of these disorders, studied through electrophysiology, show complex behaviors that go beyond simple classification as either gain or loss of function. In our study, we simulated the biophysical effects of variants (*R937C*, *V208E*, *S1336Y*, and *R853Q*) to understand their impact. Our findings reveal that all these variants negatively affect the structural stability of the gene, with *R937C* being the most unstable. Specifically, *R937C* disrupts important charged interactions affecting sodium ion flow, while *S1336Y* introduces a new interaction that impacts the channel’s inactivation gate. Conversely, the variants *V208E* and *R853Q*, which are located in the voltage-sensing domains, have opposite effects: *R853Q* increases compactness and interaction, whereas *V208E* shows a decrease. Our computer-based method offers a scalable way to gain crucial insights into how genetic variants influence channel dysfunction and contribute to neurodevelopmental disorders.

**AUTHOR SUMMARY:** Despite numerous advancements in computational methods for predicting variant pathogenicity in the *SCN2A* gene, understanding the precise biophysical molecular mechanisms associated with each variant at the atomic level remains a challenge. Presently, variants are predominantly categorized as either gain or loss of function, often overlooking critical structural details associated with these variants. This study focuses on elucidating the molecular mechanisms linked to the four most common *SCN2A* variants using all-atom molecular dynamics simulations, employing three replicas for each system. Our findings offer insights into the potential mechanisms underlying these four variants, thereby providing explanations for the observed electrophysiological outcomes. This investigation significantly contributes to enhancing our comprehension of how *SCN2A* variants manifest in various diseases. It underscores the importance of unraveling the biophysical properties underlying potential disease mechanisms, which could potentially enhance diagnostic and therapeutic strategies for patients afflicted with *SCN2A*-related disorders.

## INTRODUCTION

The sodium channel Na_V_1.2 protein, encoded by the *SCN2A* gene, belongs to the family of voltage-gated sodium ion channels (VGSCs) which are a family of integral membrane proteins that play a critical role in initiating and propagating action potentials in excitable cells(1–4). In response to membrane depolarization, VGSCs change their conformation opening the pore, allowing the passage of Na^+^ ions in the cell(1). On structure, VGSCs are heteromultimeric transmembrane proteins and consist of a 260 kDa alpha subunit which can be coupled with one or two 33-36 kDa beta subunits. These alpha subunits are large, single-chain polypeptides consisting of about 2000 amino acids and arranged in four homologous domains, namely D1 to D4. Each domain consists of six transmembrane helical segments S1 through S6. Segments S1-S4, also known as voltage-sensing domain (VSD), regulate pore opening upon membrane depolarization. The VSD domains are connected to the pore domain by a linker region between S4 and S5 segments. Segments S5-S6 of each domain come together to form a Na^+^ ion-selective pore region. This region also includes a selectivity filter (SF), the narrowest part of the pore which is formed by P-loops (the region between S5-S6 of each domain), and the inactivation gate, which is an intracellular loop connecting D-III and D-IV, which acts as a “plug” in the closing and opening of the channel. depolarization section of the pore facing the cytoplasm is formed by the combination of S5 and S6 segments from all four domains. The voltage sensors, pore region, and inactivation gate are crucial parts of the sodium channel as the majority of the pathogenic mutations in SCN genes are located in those regions(2,5,6).

Given their crucial role in brain functions, genetic variants in VGSCs genes are known to cause a wide spectrum of neurodevelopmental disorders, which can occur with or without epilepsy and have different levels of severity, onset age, and response to medication(7–9). In particular, the *SCN2A* gene is known to cause a wide range of neurodevelopmental disorders. In Clinvar, the most comprehensive database of the relationship between variants and disease phenotypes, 1839 patient variants in the *SCN2A* gene are reported, of which 449 are classified as pathogenic or likely pathogenic (accessed 12/2022). Variants in *SCN2A* have been associated with multiple disorders including autism spectrum disorder (ASD), developmental and epileptic encephalopathies (DEE), intellectual disability (ID), benign familial neonatal/infantile seizures (BFNIE), and schizophrenia(9–12). Among all VGSCs, *SCN2A* also emerges as the gene with the strongest autism spectrum disorder association(7,13,14).

At an electrophysiological level, *SCN2A* variants can modify the protein structure and its consequent ability to carry out its channel function by altering specific functional properties, such as causing a voltage shift of activation or inactivation, modifying the peak current, cell surface expression etc.(15–17). The overall net effect of these modifications caused by the variant is often summarized into one of the two functional categories of gain (GoF) and loss (LoF) of the ion channel function, depending on whether the variant enhances or reduces the overall action of the protein. These molecular effects of missense variants can determine the observed phenotype and even affect susceptibilities to pharmacological treatments. For example, in *SCN2A*-related epileptic encephalopathies, GoF missense variants are associated with an earlier seizure onset and response to sodium channel blockers. In contrast, LoF missense variants in *SCN2A* are associated with autism(18,19) and do not benefit from antiepileptic drugs(7,20). Although these molecular effects (GoF vs LoF) studied using electrophysiological experiments are crucial in identifying clinical and/or molecular phenotypes and treatments, they do not cover the entire spectrum of molecular disease mechanisms caused by a variant in the Na_V_1.2 channel(8,19) and are an over-simplification of the complex protein functionalities. For instance, if a patient has a GoF variant causing epilepsy, the sodium current might increase due to different molecular mechanisms such as a delayed closing of the inactivation gate, or a wider opening of the pore region(7,21). A recent study that investigated the functional properties of more than 30 *SCN2A* variants showed that there is a range of altered channel properties that do not fit into a simple binary GoF vs LoF classification(22). The biophysical effects behind the molecular electrophysiological heterogeneity across different missense variants is not understood.

Molecular dynamics (MD) simulations have been extensively used to capture the dynamics of molecular systems over time, offering a level of detail that is difficult to achieve through experimental techniques alone. MD simulations have proven valuable for studying various biomolecular processes, including conformational changes, protein folding, ligand-binding site prediction, and the effects of mutations at an atomic level(23,24). However, the application of MD simulations to sodium ion channels has been relatively limited, mainly due to challenges arising from their large protein structures, and limited availability of the human protein structure for all conformations. Initially, computational studies on sodium ion channels focused on only the pore region to study Na^+^ permeation, following the crystallization of the bacterial sodium channel structure(3,25),(26–28). Subsequently, with the publication of the first eukaryotic sodium channel structure, similar studies were carried out to study Na^+^ permeation pathways in eukaryotes (29,30). Focusing solely on the pore section, however, limits our ability to observe variant effects on the overall conformation of the entire protein and on other essential regions, such as the voltage-sensing domain and the inactivation gate. These regions play critical roles in the channel’s function, and understanding their structure and dynamics is crucial for a comprehensive investigation. Besides the observation of Na^+^ ion permeation, MD simulations have also been employed to understand the mechanisms of drug binding to sodium ion channels (29,31,32),(33). There has been a very limited exploration of the effects of variants in VGSCs using MD simulations. Recently, Bielopolski et al. (2022) performed virtual patch analysis via molecular dynamics simulations on six patient variants in sodium channels associated with different disease phenotypes (Dravet, epilepsy, and autism). By studying the release time of sodium ions at the DEKA motif, they predicted the functional effect of a variant that was well in agreement with existing patch clamp analysis for four out of the six variants. However, this study was limited to a relatively short simulation time of 120 ns and also did not provide information on the underlying molecular mechanism of these pathogenic variants. One of the primary reasons for the limited scope of these studies, including the use of short simulation times and the focus on the pore region, is the large size of the Na_V_1.2 protein structure, which makes MD simulations very computationally time-consuming.

In this study, we performed molecular dynamics simulations of four common Nav1.2 variants to unravel molecular disease mechanisms associated with these variants and their relationship with *in-vitro* molecular read-outs. We selected four variants (*R937C*, *V208E*, *S1336Y*, and *R853Q*) in the *SCN2A* gene based on their high prevalence in *SCN2A* patients (34,35)(Total: 22 individuals, Table 1) and on their distinct molecular read-outs, classified as either gain or loss of function based upon *in-vitro* molecular readouts. Each of the selected variants (*R937C*, *V208E*, *S1336Y*, and *R853Q*) within the *SCN2A* gene has been electrophysiologically tested and exhibits distinctive molecular characteristics and clinical phenotypes, including autism spectrum disorder (ASD) for *R937C*, the mild benign Familial Neonatal-Infantile Epilepsy (BFNIE) for *V208E*, the severe Early-Onset Epileptic Encephalopathy (EOEE) for *S1336Y*, and severe Developmental and Epileptic Encephalopathy (DEE) for *R853Q*(36–38). In this work, we carried out MD simulations for wild-type Nav1.2 protein as well as the four selected variants to uncover the disease associated mechanisms and compared this mechanism using experimental read-outs for these four variants by analyzing different biophysical properties exhibited at the atomic level. We further identified different mechanisms resulting in a loss/gain of function, providing greater detail and understanding of the variant effect.

**Table 1.** List of selected variants, their clinical annotations, and their outcomes in experimental functional tests.

| Variant | Patients | Phenotype | In vitro electrophysiological read-outs | Effect | Location |
| --- | --- | --- | --- | --- | --- |
| *R937C* | 2 | ASD | Whole cell current = 0; loss of conductance(36). | LoF | Selectivity Filter (SF) |
| *V208E* | 2 | BFNIE | Hyperpolarizing shift of activation(38). | GoF | Voltage Sensing Domain 1 (VSD-1) |
| *S1336Y* | 3 | EOEE | Smaller current density; hyperpolarized voltage dependence of activation, depolarized shift in the voltage dependence of inactivation.(37) | GoF/Mixed | Inactivation Gate (IG) |
| *R853Q* | 15 | DEE | Narrower voltage range for channel activation, slower recovery from fast inactivation, enhanced slow inactivation, hyperpolarizing shift of inactivation.<sup>22,37</sup> | LoF | Voltage Sensing Domain 2 (VSD-2) |

Phenotypes abbreviations are ASD: autism spectrum disorder; BFNIE: benign familial neonatal-infantile epilepsy; EOEE late-onset epileptic encephalopathy and d) autism spectrum disorder; DEE developmental epileptic encephalopathy.

## RESULTS

### Assessment of *SCN2A* mutations on the overall structural stability and compactness of Na_V_1.2 systems

To assess the stabilities of the Na_V_1.2 systems and examine their structural variation and dynamic alterations, we measured the root mean square deviation (RMSD) for the wild-type and each variant structure. The RMSD calculation was performed for the backbone of the protein and for a period of 500 ns (nanoseconds). We found that the average RMSD values across systems were stable after 500 ns at about ∼0.37 nm of deviation from the initial conformation. In particular, the deviations from the initial conformations for the WT and the *R937C*, *V208E*, *S1336Y*, and *R853Q* variant structures are 0.37, 0.41, 0.36, 0.32, and 0.39 nm, respectively (Figure 1a). These deviations indicate that the simulated protein-membrane complexes remained well stabilized throughout the simulation. We observed a greater RMSD change for the *R937C* variant structure at around 250 ns, when RMSD increased from ∼0.41nm to ∼0.55nm. The *R937C* variant structure exhibited the greatest deviation from its initial conformation compared to the other variant structures. Furthermore, the *R937C* structure had the highest spread of RMSD values over time. These findings indicate that all systems reached stability within the simulated time frame, and the *R937C* variant structure displays the greatest degree of conformational flexibility.

**Figure 1.**
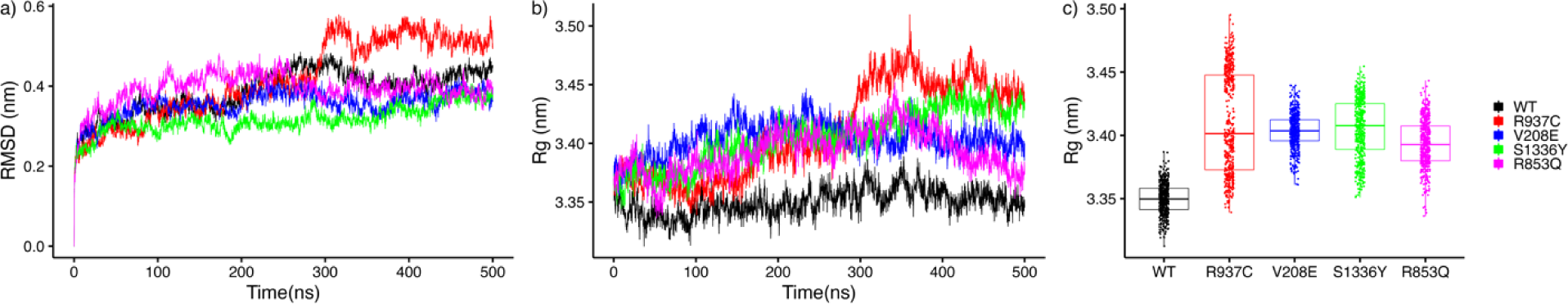
All simulated structures reach stabilization over the course of 500ns. All variant structures are less stable (a) and less compact (b and c) than the wild-type structure. The *R937C* structure is the least stable and least compact. Root-mean-square deviation (RMSD) (a) and radius of gyration (*Rg*) (b) plotted against simulation time for wild-type (WT) (black), *R937C* (red), *V208E* (blue), *S1336Y* (green), and *R853Q* (magenta) structures. c) Box plot of the radius of gyration. The colors in the legend correspond to the following variant structures: red for *R937C*, blue for *V208E*, green for *S1336Y*, and magenta for *R853Q*. Black represents the MD simulation of the wild-type structure.

We also investigated the compactness of the protein structures, which often correlates with protein compactness. To achieve this, we plotted the time-dependent radius of gyration (*Rg*) of the protein’s main chain, which is a measure of compactness, for the WT and the variant structures, using our simulation trajectory (Figure 1b). Our results revealed that all mutants underwent a structural change and were on average significantly less compact compared to the WT variant (WT*_Rg_* = 3.35 nm, *R937C_Rg_* = 3.41 nm, *V208E_Rg_* = 3.40 nm, *S1336Y_Rg_* = 3.40 nm, *R853Q_Rg_* = 3.39 nm, Wilcoxon rank sum test P < 2.2e^-16^). Across all mutants, the structure of *R937C* exhibited the widest range of compactness change across the simulation time (Figure 1c).

Notably, the increase in *Rg* after 250 ns was well-correlated with the increase in RMSD observed after 250 ns (Figure 1a), indicating structural modifications in the protein.

### Principal component analysis of the conformational space and free energy landscape

We further analyzed the conformational sampling space of wild-type (WT) and variant protein structures by examining the first two principal components (PC1 and PC2) obtained from the principal component analysis (PCA) of the positional fluctuation covariance matrix of protein Cα carbon atoms. The projection of backbone atoms onto the essential subspace, defined by principal components 1 and 2, was used to visualize the conformational sampling of the proteins. PC1 and PC2 for each protein structure were plotted at different time intervals of 100 ns each (Figure 2a). Each interval was colored according to a color gradient going from light green (time interval from 0 to 100 ns) to dark green (time interval from 400 to 500 ns) to show how the conformations evolved over time. Figure 2a shows that all variant structures explored more conformational space than WT, and all proteins reached their stable conformation after 300 ns, as evidenced by the clear overlap of conformations between the 300-400 ns interval and the 400-500 ns interval, except for the *S1336Y* structure which appeared to have multiple transient conformations.

**Figure 2.**
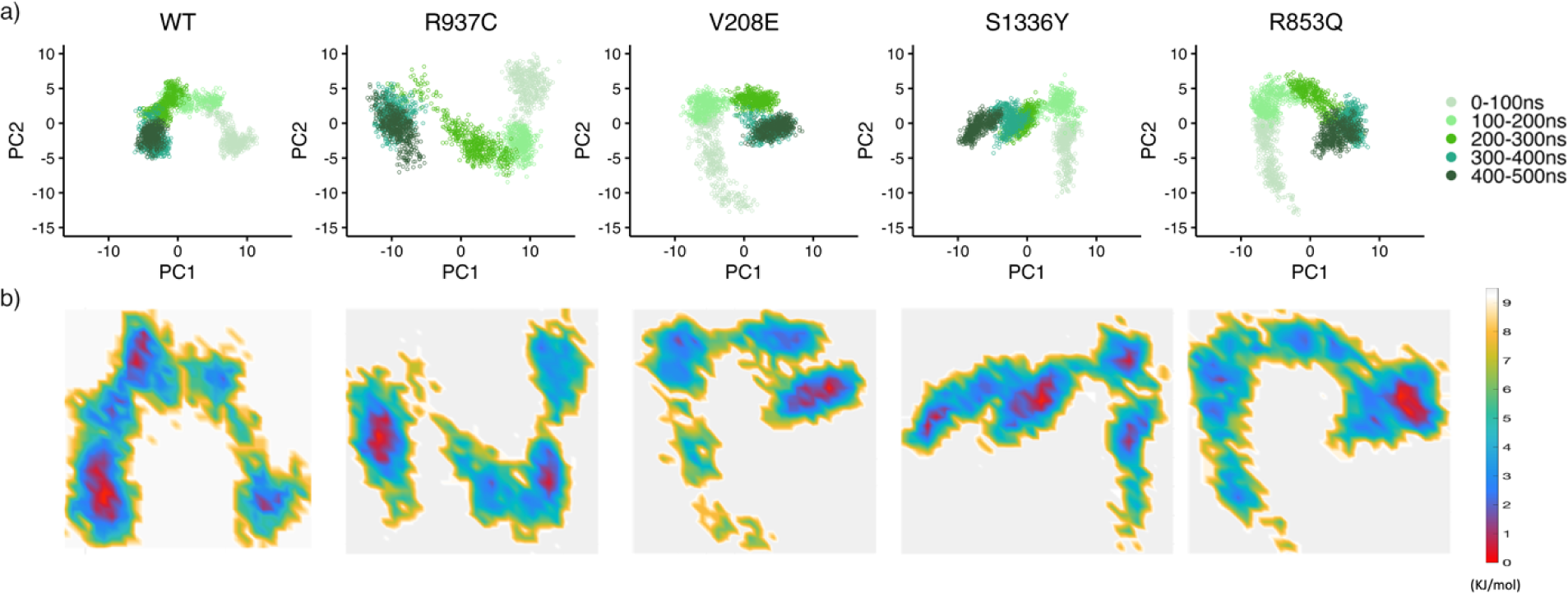
a) All variant structures explored more conformational space than WT. Projection of the Cα atoms of the WT and the four variant structures on the essential subspace, defined by the first two eigenvectors (PCs) of the covariance matrix of the wild-type protein. The colors indicate the 100 ns-long consecutive time intervals in the simulation, ranging from the lighter green color indicating a time interval from 0 to 100 ns, to the darkest green color indicating a time interval from 400 to 500 ns. b) The free energy landscape analysis shows that after 300 ns all systems stabilize on the lowest free energy conformational state. Free energy landscape analysis for WT and its variant structures. The free energy landscape was obtained using the projections of the protein’s Cα atoms position vectors onto the first two principal components as reaction coordinates. The free energy values, calculated from the distribution density, are given in kJ/mol and colored according to the color gradient detailed in the color bar. The red spots indicate the regions corresponding to the conformations with the lowest free energy and therefore more stable.

To investigate the stability of the conformational states explored by the considered structures during our simulations, we calculated the Gibbs free energy landscape using PCA, with the positional fluctuations covariance matrix of Cα atoms as reaction coordinates. The global energy minima associated with the conformational state of WT, *R937C*, *V208E*, and *R853Q*, represented by the red color of the color gradient code of Figure 2b, were achieved after 300 ns for each structure. However, for *S1336Y*, we observed multiple low free energy conformations, colored red in Figure 2b. These results suggest that although most variant structures and WT reached their stable conformation at almost the same time with minimum energy, the conformational space of mutants was more spread out before reaching their stable conformation after 300 ns.

### Pore radius

The pore region plays a crucial role in determining the selectivity and conductance properties of all ion channels. To analyze the changes introduced by the variants into the pore region, we compared the pore radius along the membrane axis in WT and variant structures, once the systems reached stability. Given that all analyzed structures reached stability after approximately 300 ns, we focused, for pore radius calculations, on the last 100 ns of the simulation in the time interval from 400 to 500 ns. We, therefore, obtained the average protein structure within the 400-500 ns time interval and utilized this structure to calculate the pore radius.

The results of the changes induced to the pore radius by the variants are shown in Figure 3. Figure 3a presents a grid surface representation of the pore radius of the WT structure that was color-coded according to radius values, where blue indicates pore regions with radius >1.6 Å, green indicates pore regions with radius comprised within 1.1 Å and 1.6 Å, and red indicates pore regions with radius <1.1 Å. Blue, therefore, indicates the most expanded regions of the pore, while red represents the most contracted ones. Figure 3b displays the grid surface representation of the pore radius for each variant structure with the same color code. Figure 3c shows the variation of the radius profile along the membrane axis for all variants when compared with WT (black line). We found that at the extracellular section of the pore, which includes the selectivity filter, the pore profiles of the variant structures exhibited minimal changes in the pore radius compared to the WT structure, with the lines in Figure 3c almost superimposed. However, more marked alterations were observed in the intracellular section of the pore. For all variant structures, we observed a reduction of the pore radius at the intracellular region in comparison to WT. The minimum value for the radius in the intracellular region is 1.38 Å for WT and 0.98 Å, 1.1 Å, 1.13 Å and 0.36 Å for *R937C*, *V208E*, *S1336Y*, and *R853Q*, respectively. These results show that the two variant structures *R853Q* and *R937C* exhibit greater pore radius contraction (Figure 3b). In these two variant structures, the observed pore contraction makes the pore radius equal (for *R937C*) and smaller (for *R853Q*) than the radius of the Na^+^ ion, which has been estimated to be 0.98 Å(51). This pore contraction to the dimension of the Na^+^ ion is likely creating a bottleneck in the pore which impedes the flow of sodium ions, subsequently causing the loss of channel function. The molecular readouts for these two variant structures indicate loss of function (LoF). Conversely, while the *V208E* and *S1336Y* variant structures also undergo pore contraction, their minimum radius (1.1 Å and 1.13 Å for *V208E* and *S1336Y*, respectively) is still greater than that of the Na^+^ ion, therefore this pore contraction is not enough to impede the passage of Na^+^ ions. Interestingly, the molecular readouts for these two variant structures indicate GoF and GoF/mixed effects.

**Figure 3.**
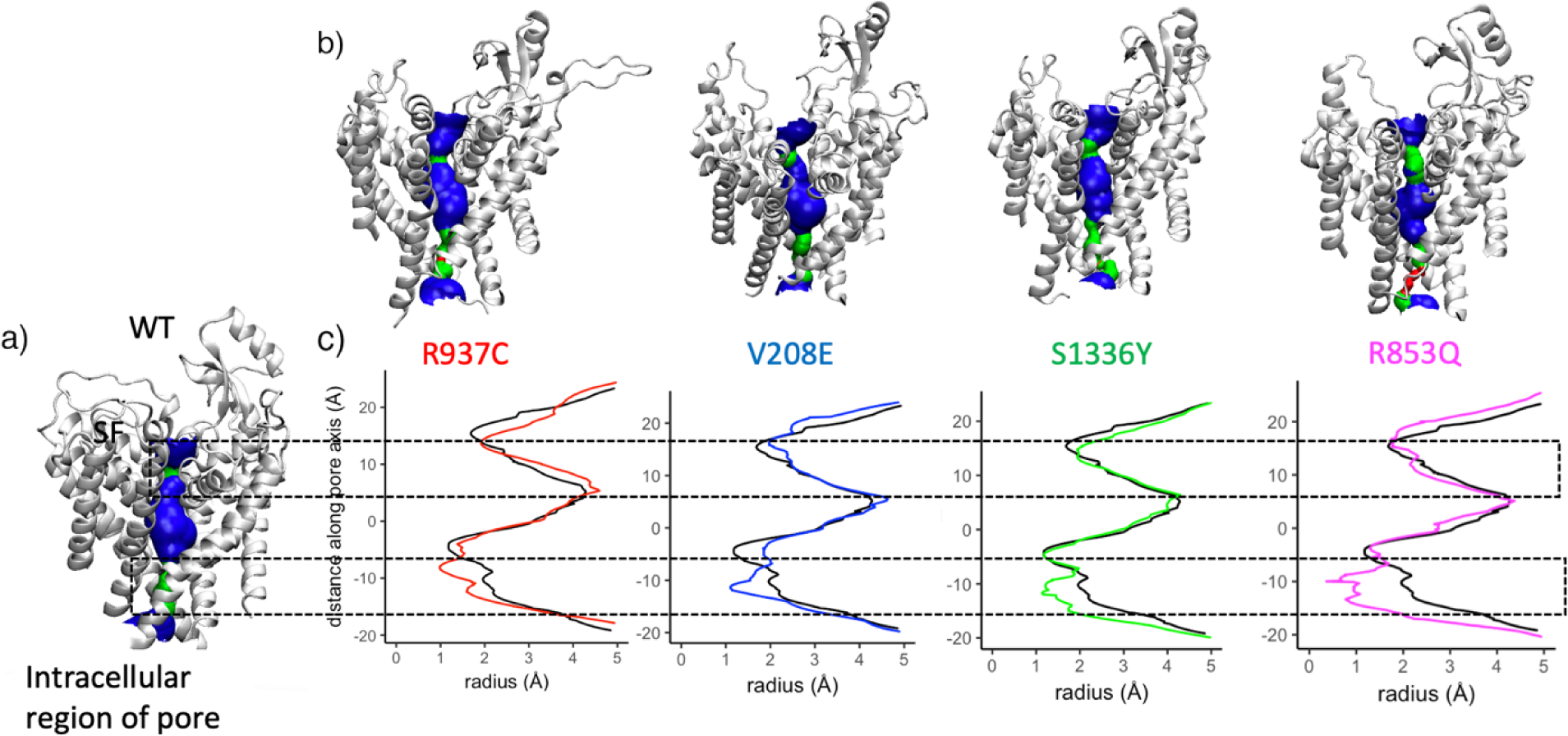
All variant structures exhibit a pore contraction in the intracellular section of the pore. The minimum radius for the *R937C* and *R853Q* variant structures becomes equal and smaller than the radius of the sodium ion (0.98 Å). In (a) and (b) is depicted the color-coded grid surface representation of the pore radius of the Nav1.2 ion channel protein, in the WT (a) and in the four (*R937C*, *V208E*, *S1336Y*, and *R853Q*) variant structures (b). Blue, green, and red indicate pore segments and are color-coded as >1.6 Å (blue), 1.1 Å-1.6 Å (green), and <1.1 Å(red). In (a) the selectivity filter and the intracellular section of the pore are highlighted by dotted lines. c) Pore radius plotted against the membrane axis. In all plots, the black line represents the WT pore radius, while the radius of the variant structures is depicted using their respective colors (red for *R937C*, blue for *V208E*, green for *S1336Y*, and magenta for *R853Q*.

### R937C disrupts two salt bridge interactions and alters the overall net charge at the selectivity filter

Upon examining the local modifications induced by the *R937C* variant, which is situated near the selectivity filter, we observed notable changes in interactions due to the substitution of arginine with cysteine. In the WT system, the positively charged R937 residue established two salt-bridge interactions with two negatively charged residues, E942 and E945. These interactions are absent in the *R937C* system (Figure 4b). To further assess the stability of these interactions, we plotted the distance from the E942 and the E945 residues of both wild-type R937 residue and variant *R937C* residue, throughout the simulation. Figure 4c illustrates the distance between *R937C*-E942 and *R937C*-E945 for both the WT (black) and variant (red) systems. The distance between both salt bridges remained stable in the WT setup, averaging approximately 3 Å apart. However, in the *R937C* setup, the distance between C937-E942 averaged around 8 Å, while the distance between C937-E945 was approximately 12 Å.

**Figure 4.**
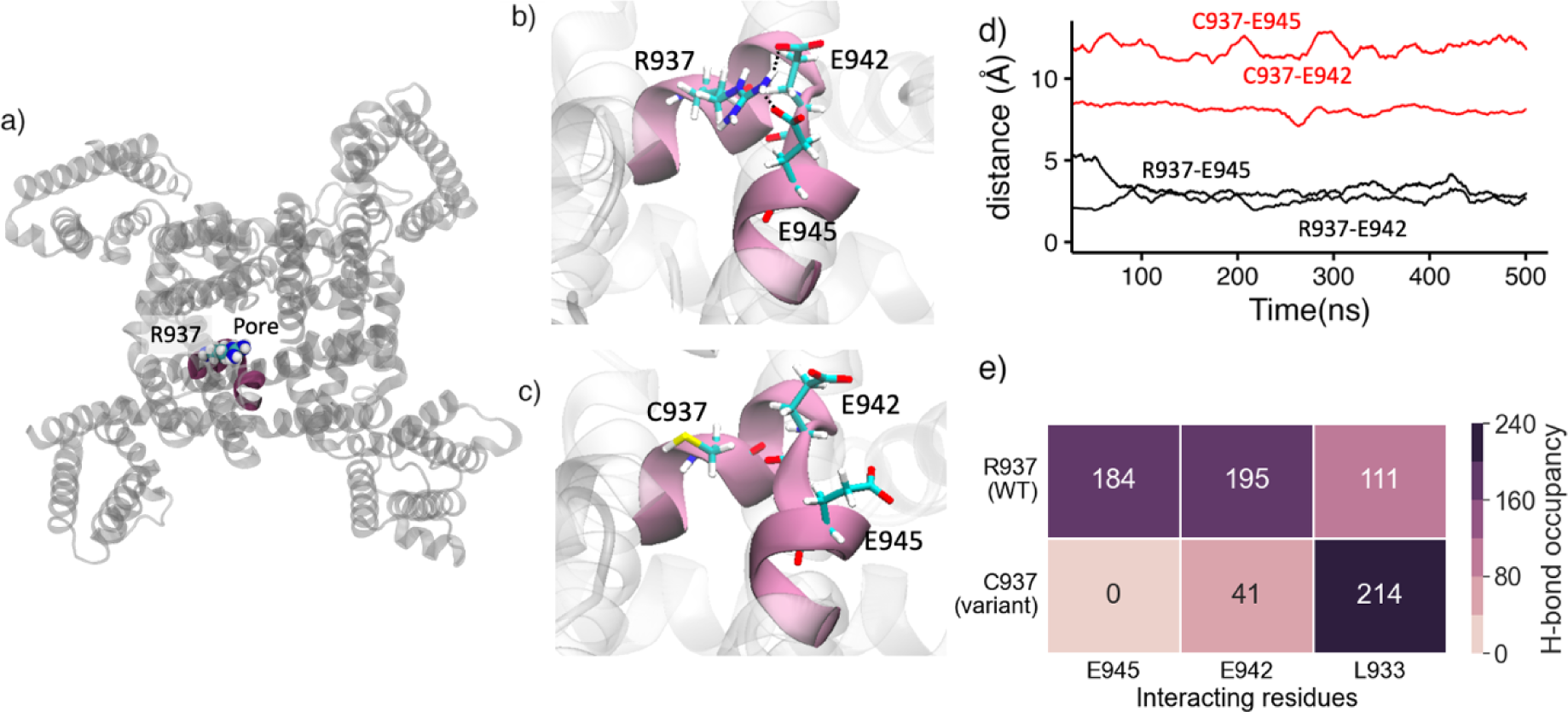
(a) Top view of the Na_V_1.2 model highlighting the position of R937 in the WT structure. (b) Close-up view showcasing the electrostatic interactions formed between R937 and E942 and E945 in WT. (c) in variant *R937C*. (d) Simulation time-dependent distance profiles of *R937C*-E942 and *R937C*-E945, represented by black (WT) and red lines (C937) respectively, illustrating the differences between the WT and *R937C*. (e) Heatmap of hydrogen bond occupancy analysis reveals the formation of hydrogen bonds by R937 and C937 with residues within a 3.5 Å cutoff distance.

To complement these findings, we also calculated the hydrogen bond occupancy between *R937C* and residues within a 3.5 Å distance from R937 in both WT and *R937C*. In MD simulation, hydrogen bond occupancy reflects the percentage of simulation time in which these interactions are formed. As depicted in Figure 4d and 6e, we observed that the WT system showed strong occupancy for the R937-E945 and R937-E942 interactions, 184% and 195%, respectively. This indicates that in the WT system, these two interactions were present during the entire simulation period of 500 ns. In contrast, in the *R937C* system, these occupancies decreased to 41% (C937-E942) and 0% (C937-E945). We also noted a change in occupancy with L933, which exhibited occupancies of 111% in the WT and 214% in the *R937C* variant. This alteration in interaction with L933 suggests that, possibly, the substitution of cysteine favors interaction with L933 rather than forming the two salt bridge interactions present in the wild-type form.

Recent electrophysiological experiments indicate that the *R937C* variant results in a complete loss of conductance, with a measured whole-cell current of 0 mA(36). Based on our findings, we propose that the absence of the two salt bridge interactions at the selectivity filter leads to a modification of the net charge at the entrance of the pore, rendering it negative. Consequently, this negative charge may hinder the movement of sodium ions through the pore, ultimately resulting in the loss of conductance in the channel. These results align well with the experimental observations.

### S1336Y forms a new interaction with the inactivation gate possibly hampering its ability to switch between the open and closed conformations

To investigate molecular mechanisms associated with GoF/mixed variant *S1336Y*, we examined the interactions formed by amino acid at position 1336 in both WT (S1336) and variant (Y1336) systems. The S1336 residue is located in domain 3 and it is part of the S4/S5 linker helix. When the serine is mutated into tyrosine at position 1336, a new hydrogen bond interaction is introduced between Y1336 and D1474, which is located in the D3-D4 connect section and is part of the inactivation gate. This interaction was quantified by plotting the distance between residue D1474 and S1336 or Y1336 for both the wild-type (WT) and variant systems, as shown in Figure 5c. The plot revealed that in the WT system, the distance between S1336 and D1474 was around 8 Å, while the distance between Y1336 and D1474 was smaller than 4 Å.

**Figure 5.**
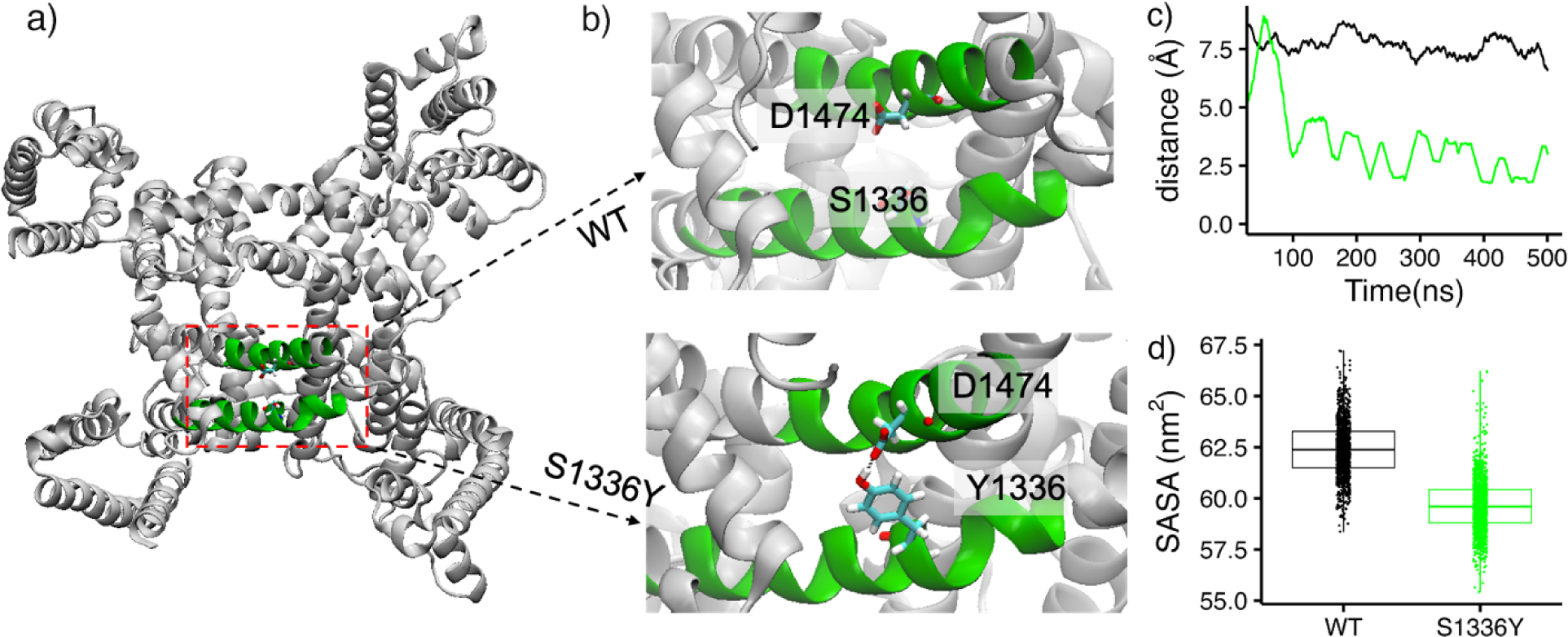
Figures (a) and (b) display the structure of the Nav1.2 protein in its wild-type form, highlighting the interaction between *S1336Y* and D1474. (c) Distance between *S1336Y* and D1474 for both the WT (in black) and *S1336Y* variant (in green) plotted against the simulation time. The plot indicates that the distance between these residues is shorter in the variant compared to the WT. (d) Box plot of the solvent accessible surface area (SASA) for the inactivation gate in both the WT (in black) and *S1336Y* variant (in green) form. The plot shows that the SASA of the inactivation gate is lower in the variant compared to the WT.

To further investigate whether this interaction changes the exposure of the inactivation gate to the solvent, the solvent-accessible surface area (SASA) for the inactivation gate was computed and plotted for both the WT and the variant, as shown in Figure 5d. The SASA plot revealed that the solvent exposure of the inactivation gate was lower in the variant (58 nm^2^) compared to the WT (62.5 nm^2^), suggesting that this new interaction stabilizes the inactivation gate. Overall, these findings suggest that a serine to tyrosine mutation at position Y1336 leads to the formation of a hydrogen bond interaction with residue D1474 which can significantly impact the structure and stability of the inactivation gate. This might also result in a modified ability of the inactivation gate to switch between open and closed conformational states, making it possible to get stuck in one particular conformation.

### Changes in compactness in opposite directions for GoF V208E and LoF *R853Q* at the voltage sensing domain

To understand the molecular mechanism associated with, GoF variant *V208E* and LoF *R853Q*, both located in voltage-sensing domains (VSD) (VSD-1 and VSD-2, respectively) we focused on structural changes and stability of these domains.

We examined the local structural flexibility and stability of these variant structures by considering the number of hydrogen bonds formed by *V208E* and *R853Q* within a 3.5 Å distance within voltage-sensing domains, as shown in Figures 8b and 8d. A distance cutoff of 3.5 Å and an angle cut-off of 30° was employed to calculate these interactions. Our results indicated that *V208E* has fewer and weaker interactions compared to the wild-type, while *R853Q* formed two new hydrogen bond interactions (Q853-L849 and Q853-S834). Next, we calculated the radius of gyration of the voltage sensing domain taking the main chain of the protein to evaluate how these interactions affect the dynamics, compactness, and stability of the voltage sensing domains. As shown in Figure 6c, VSD-2 of *R853Q* was more compact than the wild-type, suggesting increased stability. Conversely, VSD-1 of *V208E* was less compact (Figure 6a), indicating instability and suggesting a flexible voltage-sensing domain. We further performed Wilcoxon rank sum test for radius of gyration for both comparisons: WT and *V208E*; WT and *R853Q* and observed P-value of 3.59 x 10^176^ and 6.03 x 10^212^ respectively suggesting that these results are significant.

**Figure 6.**
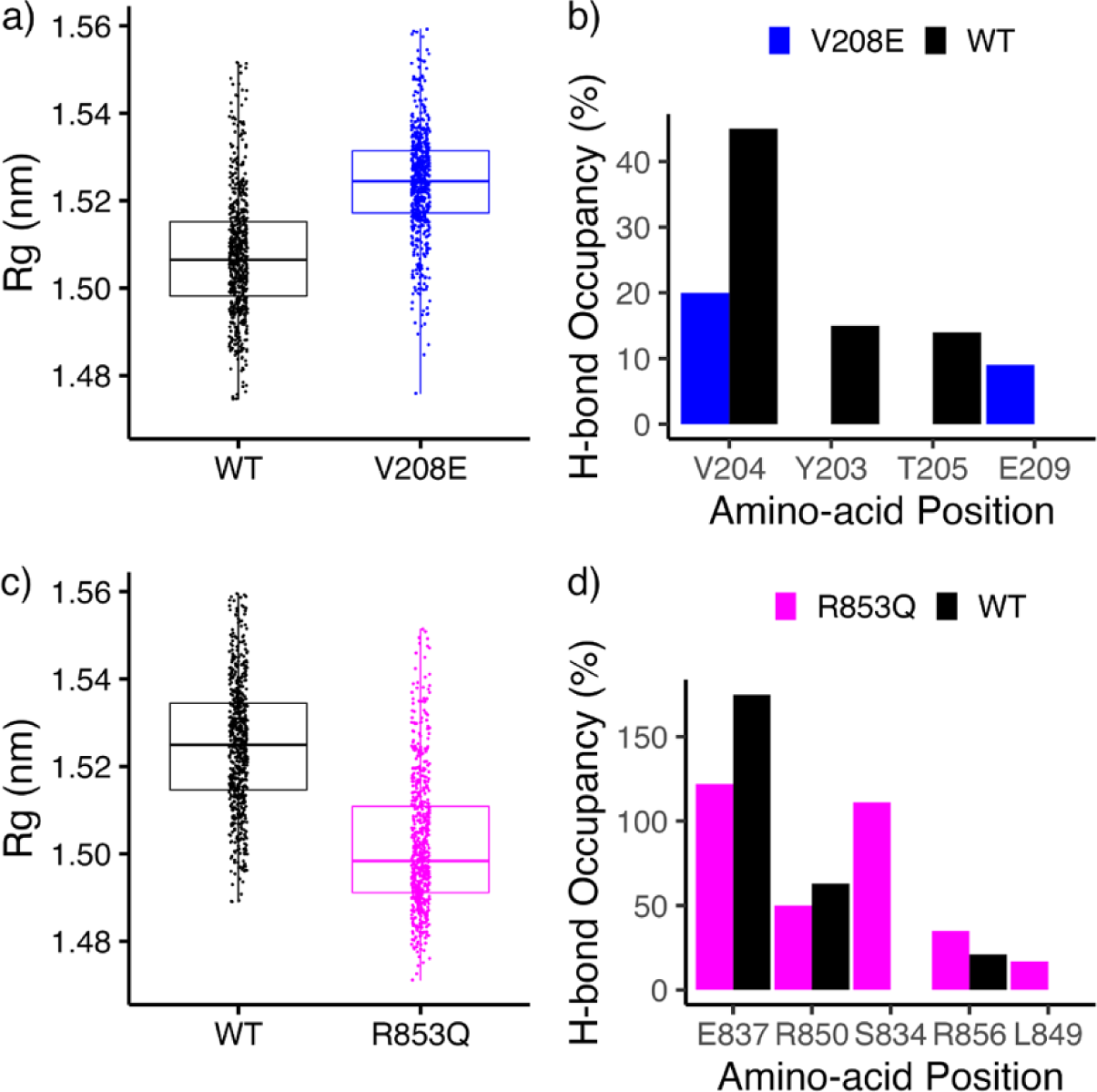
a) Box plot showing the radius of gyration (Rg) of voltage sensing domain 1 for WT (black) and *V208E* (blue). b) Hydrogen bond occupancy (%) formed between V208 and amino-acids within 3.5Å distance cutoff of V208: black (WT), blue(*V208E*) c) Radius of gyration of voltage sensing domain 2 for WT (black) and *R853Q* structure (magenta) d) Hydrogen bond occupancy (%) formed between R853 and amino-acids within 3.5Å distance cutoff of R853: black (WT), magenta (*R853Q*).

Experimental evidence demonstrates that *V208E* shifts the activation curve(38), with the hyperpolarizing effect which may facilitate easier and sooner activation of the channel thus resulting in GoF, however for *R853Q* it was observed to have a hyperpolarizing shift of inactivation(39) suggesting reduced number of channel openings, and thus, resulting in LoF. Our MD results also reveal opposite molecular mechanisms for *R853Q* and *V208E*, where *V208E* was observed to have a slightly unstable VSD-1. However, *R853Q* has more stability in VSD-2 compared to WT. This instability of VSD-1 for the *V208E* variant might help the channel to activate sooner resulting in GoF, while increased stability of VSD-2 for *R853Q* might help to stabilize the inactivated state. Our results suggest that the hyperpolarizing shift of activation observed in *V208E* may be due to the decreased stability of the VSD-1, while the hyperpolarizing shift of inactivation observed in *R853Q* may be due to the increased stability of VSD-2.

## DISCUSSIONS

Despite advancements in predicting variant pathogenicity for the *SCN2A* gene using computational methods, understanding the exact biophysical disease mechanism at an atomic level remains challenging. Current classifications often denote variants as either gain or loss, which overlooks the crucial structural information associated with these variants. To ensure effective treatment and offer informed genetic counseling, it is imperative to understand the molecular-level alterations in the channel function caused by specific genetic variations. Previous computational studies on *SCN2A* focused on understanding ion channel dynamics and very few studies performed MD simulations to understand structural changes due to *SCN2A* mutations. One such recent study performed MD simulations to predict the functional outcome of variants but was limited short simulation scale and absence of membrane. limited to a short time scale. In this study, we reveal molecular mechanisms associated with four *SCN2A* variants using *in-silico* modeling. In summary, observed changes in specific regions for each variant could affect their function. For instance, in the *R937C* variant, we identified a loss of two charged interactions, leading to a shift in the selectivity filter’s environment from neutral to negative, potentially hindering ion flow. In the *S1336Y* variant, we observed an additional interaction at the inactivation gate, possibly affecting its ability to switch between different conformations. The *V208E* and *R853Q* variants, located in voltage-sensing domains (VSD), had opposite functional effects: *V208E* displayed a gain-of-function (GoF), while *R853Q* showed a loss-of-function (LoF). Our analysis revealed structural changes in VSD for both variants, suggesting their impact on protein function. We also observed that two LoF variants (*R853Q* and *R937C*) had different molecular mechanisms, which provides a piece of evidence that two variants with the same functional effect may have different biophysical properties and possibly different disease mechanisms.

It is important to note that this study has limitations. Due to computational resource constraints, our simulations were limited to 500 ns, and although stable, longer simulations could capture additional phenomena like ion transport and gate opening/closing. Despite the considerable size of the sodium ion channel protein, we conducted three replicas for each system, each lasting 500 ns for each variant focusing on the transmembrane region of the protein, which was not done in prior MD studies.

Despite some limitations, our findings provide insights into the potential mechanisms for four variants, providing explanations for the observed electrophysiological outcomes. This study contributes to a better understanding of how *SCN2A* variants lead to different diseases, emphasizing the elucidation of biophysical properties that underlie possible disease mechanisms, and potentially aid in improving diagnosis and treatment approaches for patients with *SCN2A*-related disorders.

## METHODS

### Variant selection

Based on these three criteria, we identified four missense variants: *V208E*, *R853Q*, *R937C*, and *S1336Y* each associated with a different functional outcome, which we incorporated into our study and are described in Table 1. Our selection was based on three criteria. First, the variant must have been experimentally functionally tested, with literature having reached agreement on its either GoF or LoF effect. Second, the variant must have been detected in multiple patients and these patients should exhibit diverse clinical phenotypes. This second requirement stems from the observations of correlations between specific clinical phenotypes and protein functions (LoF/GoF/Mixed)(19), therefore the associated phenotypes could provide supporting evidence for the different molecular changes of the selected variants. The selected variants should represent different phenotypes as represented in Table 1. Third and last criterion, in alignment with the suggestions from past research that the functional impacts of these variants may correlate with their three-dimensional localization within the Na_V_1.2 protein structure, we ensured that the selected variants were located at different protein domains.

### Modeling of the Na_V_1.2 protein structure

In this work, our goal was to simulate the Na_V_1.2 protein in the open conformation to study its dynamics in the wild-type (WT) and in the four mutants. As there are no resolved structures of Na_V_1.2 in the open conformation, we used homology modeling to model the Na_V_1.2 structure.(31), onto a recently crystallized open conformation structure of SCN5A gene protein, Na_V_1.5(40). Homology modeling is a valid approach given the significant sequence similarity observed among human Na_V_ channels in the transmembrane region, which ranges from 85% to 91%. This high level of similarity ensures a consistent and reliable output from the comparative modeling process(41). We performed homology modeling using *Swiss model* application and structurally aligned the modeled Na_V_1.2 and the crystal structure of Na_V_1.2 (PDB-ID: 6j8e). The aligned structures have a RMSD value of 0.98 Å, indicative of a good alignment. We did notice a slight difference in conformation between the structures at the inactivation gate as shown by the dotted circle in Figure 7b. Moving forward, we used the modeled Na_V_1.2 structure as the initial structure for all our MD simulations.

**Figure 7.**
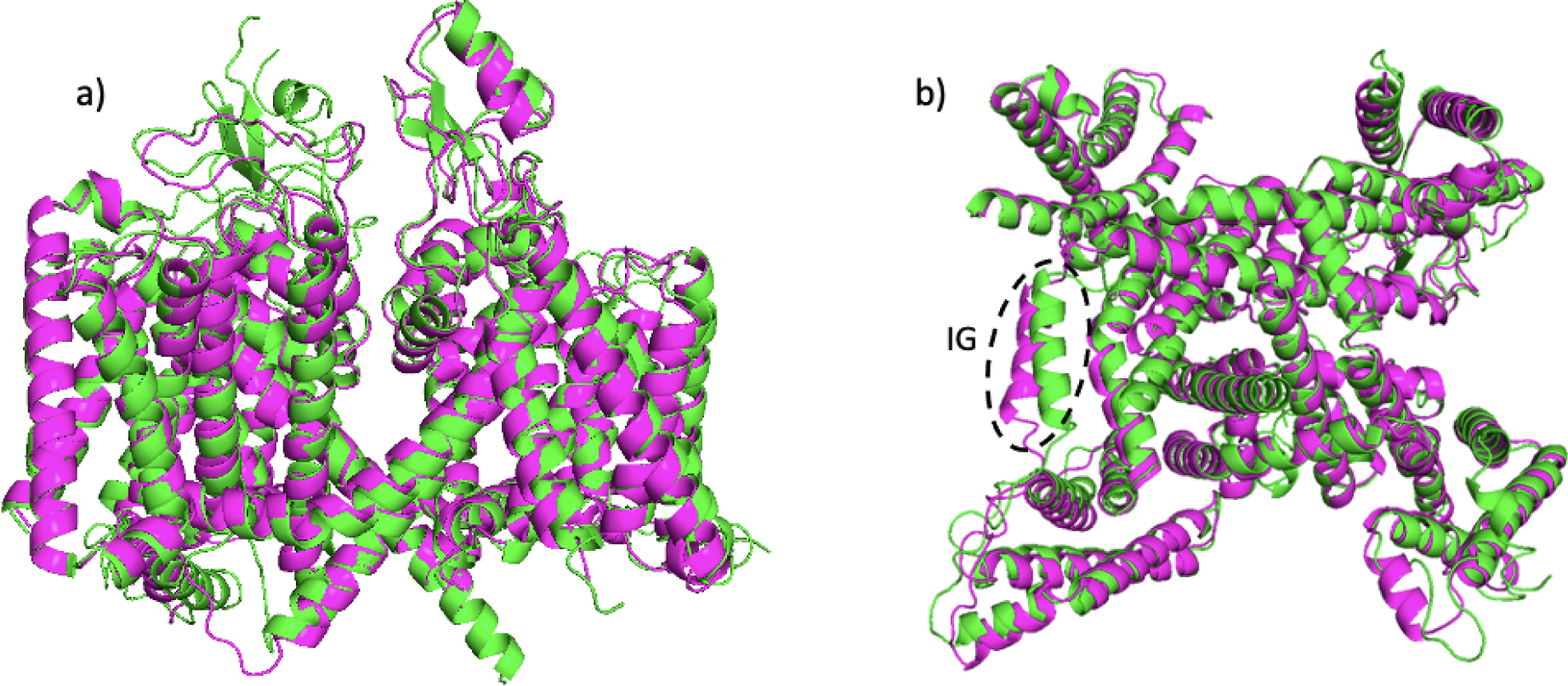
Structural alignment of the homology modeled structure of Na_V_1.2 protein (green) on the Na_V_1.2 protein structure (PDB-ID: 6j8e, magenta). **(a)** Side view; **(b)** Top view where dotted region represents the difference in inactivation gate region conformation

### Protein-membrane setup

The setup for our MD simulations was done according to the standard protocol for protein-membrane systems, as follows. We constructed five Na_V_1.2 protein membrane models (WT and four mutants) using the Charmm-GUI web server’s membrane builder(42) application. Only the transmembrane region, including the pore and voltage sensing domains, was considered for system construction. The PPM(positioning of proteins in membranes) server within Charmm-GUI was used to predict the protein’s membrane-interacting region. The resulting protein-membrane systems consisted of POPC (1-palmitoyl-2-oleoyl-sn-glycero-3-phosphocholine) lipid molecules, with dimensions of 185 x185 x186 Å^3^, and were solvated with TIP3 water molecules. 0.15 M NaCl ions were added to neutralize the system. After system assembly, we performed all-atom molecular dynamics simulations using Gromacs 2021 software(43) and charmm36 forcefield(44). The steepest descent algorithm was used for energy minimization of each system for 5000 steps. To equilibrate the systems, we followed six consecutive equilibration steps, gradually decreasing the harmonic restraint for proteins and lipids. The equilibration steps are summarized in (Supplementary Table 1), with the first two steps carried out in the NVT (constant volume, temperature, and particle number) ensemble and the remaining four steps in the NPT (constant pressure, temperature, and particle number) isobaric-isothermal ensemble.(45)

### Gromacs simulation protocol

After performing the energy minimization and the equilibration steps, we carried out isobaric-isothermal (NPT) production runs for each system using 2 fs steps, employing semi-isotropic Parrinello-Rahman(46) pressure coupling at a pressure of 1 bar. To constrain the bonds involving hydrogen atoms, we used the LINCS(47) algorithm. Nose-Hoover(45) temperature coupling was used to maintain the temperature of the models. The van der Waals and electrostatic interactions were truncated at a cutoff of 12 Å, and the Particle Mesh Ewald (PME)(48) method was used to handle long-range electrostatic interactions with periodic boundary conditions. A representative initial configuration for the WT system is presented in Figure 8. To ensure the reproducibility and reliability of our results, we performed two additional replicas of each system for 500 ns.

**Figure 8.**
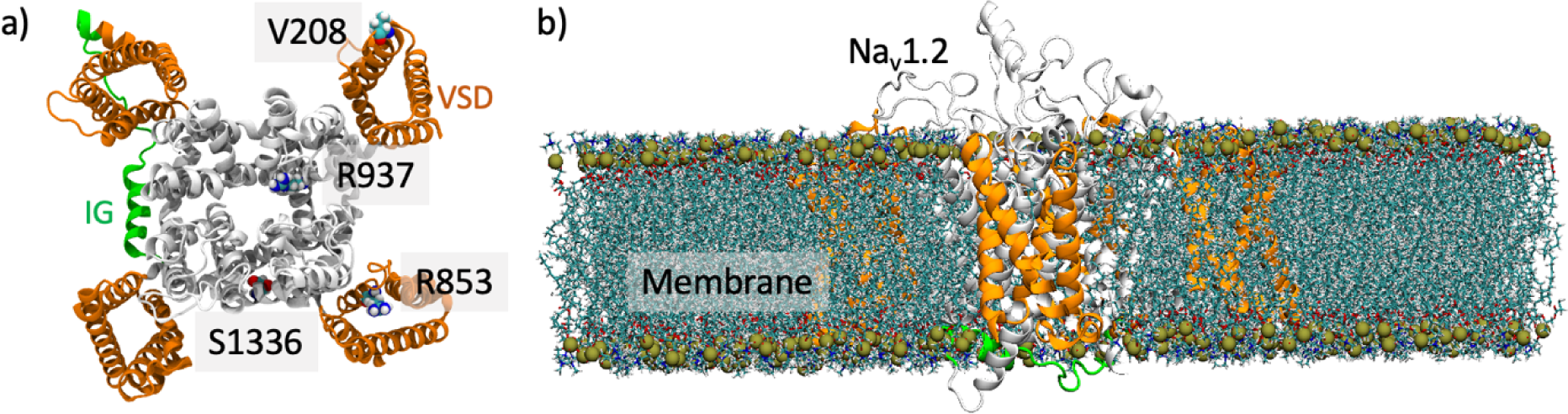
a) Top view of the Na_V_1.2 model. Initial WT protein conformation with the four voltage-sensing domains (VSD) and the inactivation gate (IG) colored in orange and green, respectively. The four residues whose variants are considered in this study are highlighted through sphere representation. b) Side view of the WT protein-membrane set up at 0ns.

### Analysis of MD trajectories

All the analyses were carried out using embedded Gromacs modules in the Gromacs simulation package and Visual Molecular Dynamics (VMD). We analyzed our files using *gmx_rms, gmx_rmsf, gmx_sham, gmx_gyrate, gmx_anaeig, gmx_covar, gmx_sasa* to extract root mean square deviation, root mean square fluctuations, radius of gyration, solvent accessible surface area, etc. All the graphs were plotted using R (version 4.2.0). Visualizations were done using VMD software (version 1.9.4)(49). The pore radius was calculated using the *hole* program with default settings(50).

### 2.6. Principal components analysis (PCA) and Free energy landscape (FEL)

We calculated principal component analysis (PCA) of Cα atoms for WT and mutants. We used the *gmx_covar* module which performs calculation and diagonalization of the covariance matrix and outputs the corresponding eigenvectors and eigenvalues based upon positional fluctuations of C_α_ atoms as described as *C*_*ij*_= 〈 (*x*_*i*_ ― 〈*x*_*i*_〉) (*x*_*ij*_― 〈*x*_*j*_〉) 〉 where x_i_/x_j_ is the coordinate of i/j_th_ atom and < > represents ensemble average.

We calculated the Free Energy Landscape (FEL), which is useful for understanding the stability, folding, and extracting minimal energy conformation of a protein. The FEL can be constructed using the following equation: *G*(*x*) = ― *k*_*B*_*T*ln *P*(*x*) here, *k*_*B*_ represents the Boltzmann constant, *T* represents the absolute temperature, and *P*(*x*) represents the probability distribution of the molecular system along the principal components (PCs).

## Acknowledgment

I.N.S. is funded, in part, by the Ambrose Monell Cancer Genomic Medicine Fellowship and the NIH National Institute of General Medical Sciences (NIGMS) Maximizing Opportunities for Scientific and Academic Independent Careers (MOSAIC) K99/R00 grant – 1K99GM143552.

